# PelX is a UDP-*N*-acetylglucosamine C4-epimerase involved in Pel polysaccharide-dependent biofilm formation

**DOI:** 10.1101/2020.05.26.111013

**Authors:** Lindsey S. Marmont, Gregory B. Whitfield, Roland Pfoh, Rohan J. Williams, Trevor E. Randall, Alexandra Ostaszewski, Erum Razvi, Ryan A. Groves, Howard Robinson, Mark Nitz, Matthew R. Parsek, Ian A. Lewis, John C. Whitney, Joe J. Harrison, P. Lynne Howell

**Affiliations:** Program in Molecular Medicine, The Hospital for Sick Children, Toronto, ON, Canada; Department of Biochemistry, University of Toronto, Toronto, ON, Canada; Department of Chemistry, University of Toronto, Toronto, ON, Canada; Department of Biological Sciences, University of Calgary, Calgary, AB, Canada; Photon Science Division, Brookhaven National Laboratory, Upton, NY, USA; Department of Microbiology, University of Washington, Seattle, WA, USA

**Author notes:** Department of Microbiology, Harvard Medical School, Boston, MA, USA. Département de microbiologie, Infectiologie et Immunologie, Université de Montréal, Montréal, Quebec, Canada (Gregory B. Whitfield). Department of Biochemistry and Biomedical Sciences, McMaster University, Hamilton, ON, Canada. These authors contributed equally to this work (Lindsey S. Marmont, Gregory B. Whitfield). Address correspondence to: P. Lynne Howell.

**Keywords:** polysaccharide, biofilm, *Pseudomonas protegens*, X-ray crystallography, epimerase

## Abstract

Pel is an *N*-acetylgalactosamine rich polysaccharide that contributes to the structure and function of *Pseudomonas aeruginosa* biofilms. The *pelABCDEFG* operon is highly conserved among diverse bacterial species, and thus Pel may be a widespread biofilm determinant. Previous annotation of *pel* gene clusters led us to identify an additional gene, *pelX*, that is found adjacent to *pelABCDEFG* in over 100 different bacterial species. The *pelX* gene is predicted to encode a member of the short-chain dehydrogenase/reductase (SDR) superfamily of enzymes, but its potential role in Pel-dependent biofilm formation is unknown. Herein, we have used *Pseudomonas protegens* Pf-5 as a model to understand PelX function as *P. aeruginosa* lacks a *pelX* homologue in its *pel* gene cluster. We find that *P. protegens* forms Pel-dependent biofilms, however, despite expression of *pelX* under these conditions, biofilm formation was unaffected in a Δ*pelX* strain. This observation led to our identification of the *pelX* paralogue, PFL_5533, which we designate *pgnE*, that appears to be functionally redundant to *pelX*. In line with this, a Δ*pelX* Δ*pgnE* double mutant was substantially impaired in its ability to form Pel-dependent biofilms. To understand the molecular basis for this observation, we determined the structure of PelX to 2.1Å resolution. The structure revealed that PelX resembles UDP-*N*-acetylglucosamine (UDP-GlcNAc) C4-epimerases and, using ^1^H NMR analysis, we show that PelX catalyzes the epimerization between UDP-GlcNAc and UDP-GalNAc. Taken together, our results demonstrate that Pel-dependent biofilm formation requires a UDP-GlcNAc C4-epimerase that generates the UDP-GalNAc precursors required by the Pel synthase machinery for polymer production.

## INTRODUCTION

Exopolysaccharides are a critical component of bacterial biofilms. The opportunistic pathogen *Pseudomonas aeruginosa* is a model bacterium for studying the contribution of exopolysaccharides to biofilm architecture because biofilms formed by this organism use exopolysaccharides as a structural scaffold (1). *P. aeruginosa* synthesizes the exopolysaccharides alginate, Psl, and Pel, and each have been shown to contribute structural and protective properties to the biofilm matrix under various conditions (2). While these polysaccharides differ in their chemical composition and net charge, the synthesis of all three polymers requires sugar-nucleotide precursors. Genes encoding enzymes required for precursor generation are often found within or adjacent to the gene cluster responsible for the production of their associated polysaccharide. For example, Psl requires GDP-mannose (GDP-Man) precursors, which are generated from mannose-1-phosphate by the enzyme PslB (3). Similarly, alginate requires the precursor GDP-mannuronic acid (GDP-ManUA) and the *alg* locus encodes two of the three enzymes, AlgA and AlgD, required to synthesize this activated sugar (4,5). The third enzyme, AlgC, is not found within the *alg* operon and is also involved in synthesizing precursors for Psl and B-band lipopolysaccharide (6).

In Gram-negative bacteria, the *pelABCDEFG* operon encodes seven gene products that are required for <u>pel</u>licle (Pel) biofilm formation (7). These biofilms form at the air-liquid interface of standing *P. aeruginosa* cultures (8). In contrast to the Psl and alginate gene clusters, none of the *P. aeruginosa pel* genes are predicted to be involved in sugar-nucleotide precursor production, indicating that, like AlgC, these functions are encoded by genes elsewhere on the chromosome. Analyses of Pel have demonstrated that it is a cationic polysaccharide rich in *N*-acetylgalactosamine (GalNAc) residues and that the putative Pel polymerase, PelF, preferentially interacts with the nucleotide UDP (9). Additionally, functional characterization of PelA has demonstrated that it is a bifunctional enzyme with both polysaccharide deacetylase and α-1,4-*N*-acetylgalactosaminidase activities, which further supports the hypothesis that the precursor required for the biosynthesis of Pel is an acetylated sugar (10,11). Together, these data suggest that a key sugar-nucleotide precursor involved in Pel biosynthesis is UDP-GalNAc, the high energy precursor needed for the biosynthesis of GalNAc-containing glycans.

We recently made the observation that many bacteria possess an additional open reading frame in their *pel* biosynthetic gene clusters that is predicted to encode a member of the short-chain dehydrogenase/reductase (SDR) enzyme superfamily (12,13). The SDR superfamily is an ancient enzyme family whose members share a common structural architecture and are involved in the synthesis of numerous metabolites, including sugar-nucleotide precursors used for the generation of bacterial cell surface glycans (14). In several species of bacteria, such as the plant-protective Pseudomonad *P. protegens*, the SDR encoding gene *pelX* is found directly upstream of the *pel* genes while other bacteria such as *P. aeruginosa* lack this gene within their *pel* locus.

Plant root colonization by *P. protegens* Pf-5 requires the formation of biofilms. This process has been shown to require the biofilm adhesin, LapA (15). In addition to LapA, biofilms produced by this strain also contain undefined exopolysaccharides (16,17). Besides Pel, *P. protegens* Pf-5 has the genetic capacity to synthesize the exopolysaccharides Psl, alginate, and poly-β-1,6-*N*-acetylglucosamine (PNAG), however, little is known about the role these polymers play in *P. protegens* biofilm formation (17).

In the present study, we show that *P. protegens* Pf-5 forms Pel-dependent biofilms at air-liquid interfaces and using *P. protegens* PelX as a representative Pel polysaccharide-linked SDR enzyme, we find that this enzyme functions as a UDP-GlcNAc C4-epimerase. We find that the *pelX* gene is not essential for Pel-polysaccharide-dependent biofilm formation because *P. protegens* possesses a paralogue of this gene, PFL_5533. Deletion of both of these genes was found to substantially impair Pel-dependent biofilm formation. Based on our analyses we designate PFL_5533 as <u>p</u>olysaccharide UDP-<u>G</u>lc<u>N</u>Ac <u>e</u>pimerase (*pgnE*) and propose that the production of UDP-GalNAc by UDP-GlcNAc C4-epimerases is a critical step in the biosynthesis of the Pel polysaccharide.

## RESULTS

### *Identification of a SDR family enzyme associated with* pel *gene clusters*

In a previous study, we used the sequence of PelC, a protein required for Pel polysaccharide export, to identify *pel* biosynthetic loci in a wide range of Proteobacteria (12). In addition to the conserved *pelABCDEFG* genes, several of these loci contained an additional open reading frame. We observed several genomic arrangements containing this gene **(Fig. 1)**. In 70% of these genomes, the additional gene is located directly upstream of *pelA* and may be transcribed together with the *pel* genes. In 24% of cases, the gene is located upstream of *pelA* but is divergently transcribed, while 5% of the time the gene is encoded downstream of *pelG* **(Fig. 1)**. Sequence and structure-based analyses of the protein product of this gene, PelX, using BLAST and Phyre^2^ suggest that it likely encodes an SDR family enzyme (18,19). In total, we identified 136 *pel* loci containing a *pelX* gene **(Fig. 1, Data Set S1)**.

**Figure 1:**
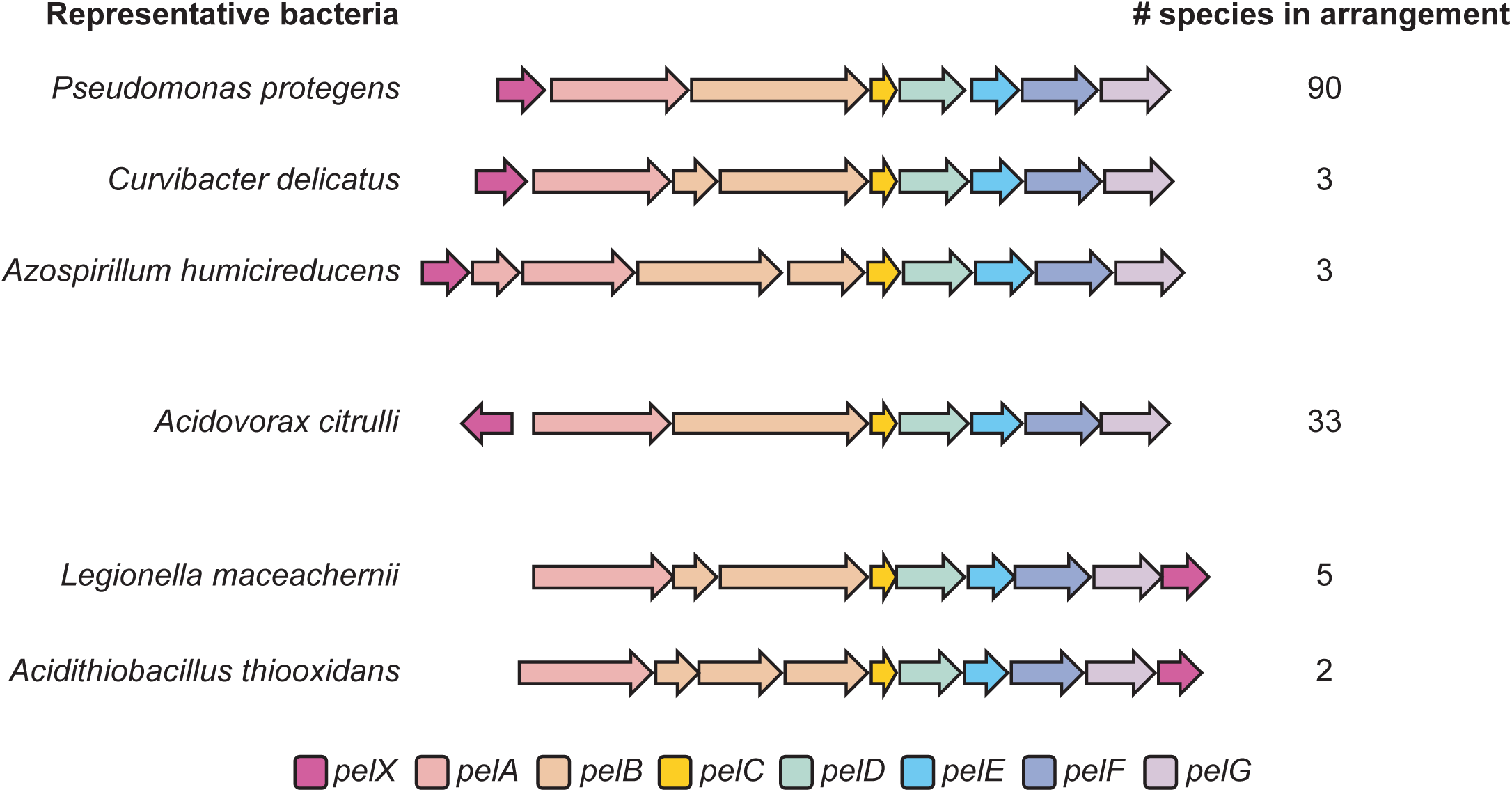
A *pelX* homologous gene is found adjacent to *pel* biosynthetic gene clusters in three different arrangements. Each gene is shown as an arrow, where each arrow indicates the direction of transcription. The predicted function of each gene is indicated by its colour as per the legend (bottom). One representative bacterial species is shown per gene arrangement. The total number of species identified with each arrangement is indicated (right). The full list of species with *pelX*-containing *pel* loci can be found in **Dataset S1**.

In order to determine whether *pelX* plays a role in Pel polysaccharide dependent biofilm formation, we set out to characterize *pelX* in a species of bacteria for which the regulation of *pel* gene expression has been studied. In *P. protegens*, which contains a *pelX* gene upstream of *pelA*, the *pel* gene cluster is under the control of the same Gac/Rsm global regulatory cascade as in *P. aeruginosa* (17). In addition, two putative recognition sequences for the enhancer binding protein FleQ are found upstream of *pelX* (PFL_2971), not *pelA*, suggesting that in contrast to *P. aeruginosa, pelX* may be the first gene of the *pel* operon in this species (20). Given that this operon is likely regulated in a similar manner to the *pel* locus of *P. aeruginosa* and that these two species are closely related, we used *P. protegens* to characterize the role of PelX in biofilm formation.

### P. protegens forms Pel-dependent biofilms that are enhanced by elevated levels of c-di-GMP

In addition to the *pel* genes, *psl* gene expression has been shown to be regulated by the Gac/Rsm pathway in *P. protegens* and this regulatory cascade is required for *P. protegens* biofilm formation (17). Interestingly, some strains of *P. aeruginosa*, including PAO1, use Psl as their predominant biofilm matrix exopolysaccharide whereas others, such as PA14, use Pel (21). Therefore, in order to determine whether *P. protegens* biofilms are dependent on Pel and/or Psl, we generated strains lacking *pelF* or *pslA*, genes previously shown to be required for Pel- and Psl-dependent biofilm formation, respectively, and examined whether these strains could form biofilms (8,22). After five days of static growth in liquid culture, we found that wild-type and Δ*pslA* strains of *P. protegens* adhered similarly to a polystyrene surface, whereas a strain lacking *pelF* displayed a marked reduction in surface attachment **(Fig. 2A)**. The level of surface adherence of a Δ*pelF* Δ*pslA* double mutant was comparable to that of the Δ*pelF* strain. Based on these data, we conclude that the Pel polysaccharide is a critical component of *P. protegens* Pf-5 biofilms.

**Figure 2:**
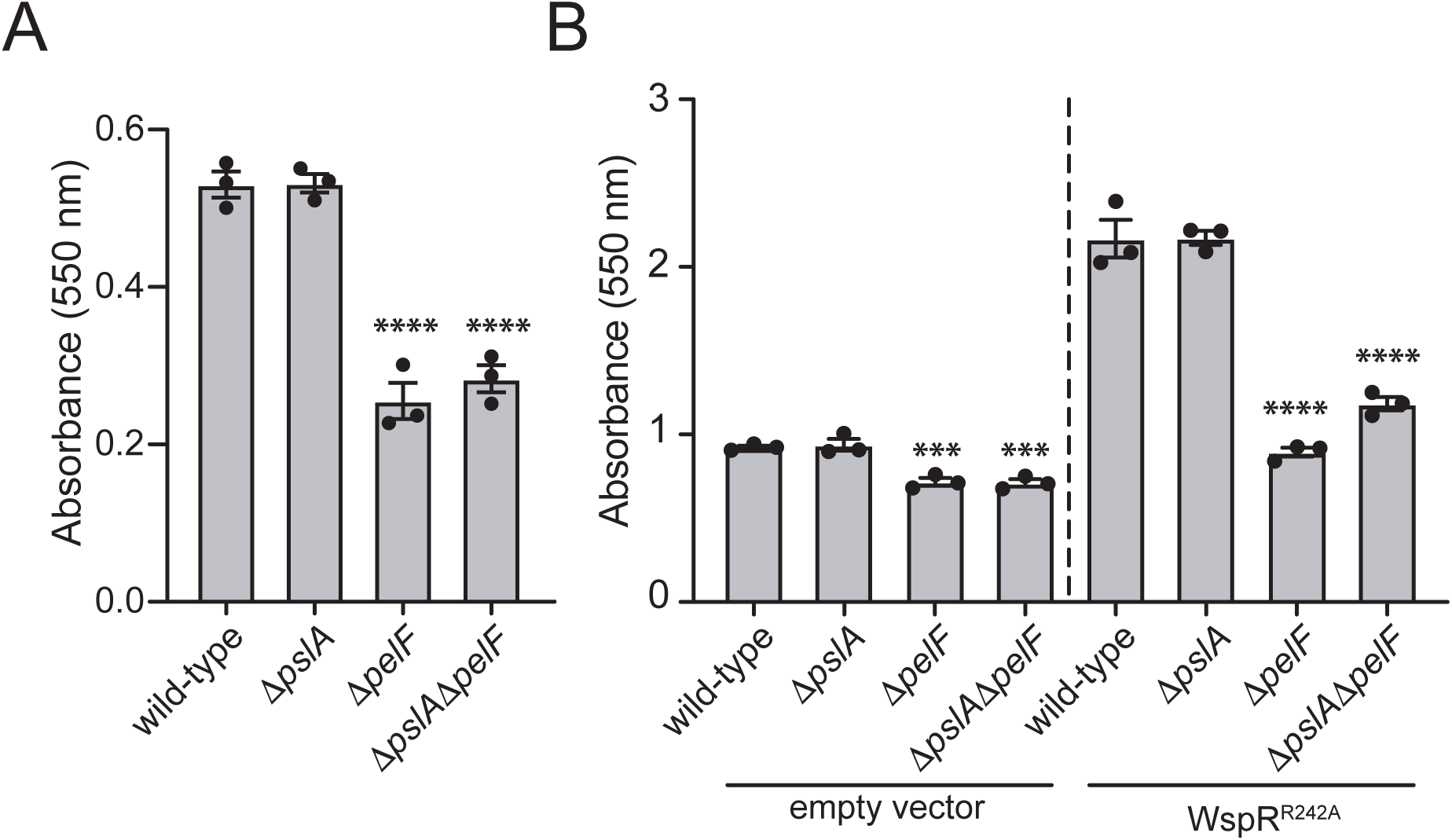
*P. protegens* forms Pel polysaccharide-dependent biofilms that are enhanced by the overexpression of the diguanylate cyclase, WspR. Biofilm biomass determined using the crystal violet assay for (A) wild-type *P. protegens* Pf-5 and the corresponding deletion mutants assayed after 5 days of growth, or (B) strains expressing pPSV39 (empty vector) or pPSV39::*wspR*^*R242A*^ after 24 h of growth. Error bars indicate the standard error of the mean of two (A) or three (B) independent trials performed in triplicate. Statistical significance was evaluated using one-way ANOVA with Bonferroni correction; ***P < 0.001; ****P < 0.0001.

Previous analysis of the region upstream of *P. protegens pelX* identified a FleQ consensus binding sequence (20). FleQ is a bis-(3′,5′)-cyclic dimeric guanosine mono-phosphate (c-di-GMP) responsive transcription factor that binds to specific sequences upstream of the *pel* operon in *P. aeruginosa*, blocking their transcription (23). When the intracellular concentration of c-di-GMP is high, FleQ switches to an activator and upregulates transcription of the *pel* genes (24). Based on these observations, we reasoned that expression of the *P. protegens pel* operon is likely upregulated in the presence of elevated levels of c-di-GMP (23). To test this hypothesis, we expressed the well-characterized diguanylate cyclase WspR of *P. aeruginosa* from an IPTG-inducible plasmid in *P. protegens* (25). Because WspR activity can be inhibited by c-di-GMP binding to an allosteric site of the enzyme, we inactivated this autoinhibitory site by introducing a previously characterized R242A point mutation into the sequence of the protein (WspR^R242A^; (26)). Upon induction of WspR^R242A^ expression, approximately 2.3-fold more *P. protegens* adhered to polystyrene surfaces compared to a vector control strain **(Fig. 2B)**. Taken together, our data suggests that *P. protegens* Pel-dependent biofilm formation is enhanced in response to elevated intracellular c-di-GMP levels.

### pelX *is expressed under biofilm promoting conditions but is functionally redundant with PFL_5533*

Since Pel-dependent biofilm formation is enhanced in the presence of c-di-GMP, and FleQ is predicted to bind upstream of the *pelX* gene, we reasoned that *pelX* is most likely expressed in a c-di-GMP dependent manner along with the rest of the *pel* genes. To test this, we probed for the expression of PelX by fusing a vesicular stomatitis virus glycoprotein (VSV-G) tag to its C-terminus at the native *pelX* locus on the *P. protegens* chromosome. To examine expression of the *pel* operon, a VSV-G tag was similarly added to the C-terminus of the putative Pel synthase subunit, PelF (13). Strains expressing either WspR^R242A^ or a vector control were grown under biofilm-conducive conditions and analyzed by Western blot. In strains lacking WspR^R242A^, neither PelX nor PelF could be detected; however, in the WspR^R242A^ expressing strains, both PelX and PelF were detected at their expected molecular weights of 34 and 58 kDa, respectively **(Fig. 3A).** These data suggest that *pelX* and *pelF* expression are positively regulated by c-di-GMP in *P. protegens*, and that PelX is expressed under conditions where the Pel polysaccharide is produced. However, when we deleted *pelX*, we found that *P. protegens* biofilm biomass was unaffected, indicating that PelX is not essential for Pel-dependent biofilm formation **(Fig. 3B)**. These findings led us to hypothesize that the *P. protegens* genome might encode a second SDR enzyme that renders PelX functionally redundant. We queried the PelX amino acid sequence against the *P. protegens* Pf-5 proteome using BLASTP to identify similar proteins (18). This search identified several proteins from the SDR superfamily (**Table 1**), however, one protein in particular, PFL_5533, stood out because it shares 68% sequence identity with PelX. To determine whether PFL_5533 is expressed during *P. protegens* biofilm formation, we fused a C-terminal VSV-G tag to PFL_5533 at its native chromosomal locus and examined its expression in the presence and absence of WspR^R242A^. We detected similar levels of VSV-G tagged PFL_5533 in both vector control and WspR^R242A^-expressing strains suggesting that in contrast to *pelX*, the expression of this gene does not change in response to c-di-GMP **(Fig. 3A).** The observation that PFL_5533 is expressed during biofilm growth conditions and that it possesses high sequence homology to *pelX* led us to probe its potential role in Pel polysaccharide production.

**Table 1:** Short-chain dehydrogenase/reductase enzymes in *P. protegens* Pf-5.

| Pf-5 PFL number | Annotation | Predicted function<sup>a</sup> | % Identity to PelX |
| --- | --- | --- | --- |
| PFL_2971 | PelX | NAD-dependent epimerase/dehydratase | 100.00 |
| PFL_5533 |  | NAD-dependent epimerase/dehydratase | 67.97 |
| PFL_3079 | LspL | UDP-glucuronate 5'-epimerase | 30.54 |
| PFL_5405 | GalE | UDP-Glc 4-epimerase | 29.88 |
| PFL_0305 | rfbB | dTDP-glucose 4,6-dehydratase | 33.60 |
| PFL_5490 |  | NAD dependent epimerase/dehydratase | 30.87 |
| PFL_4822 |  | 3-beta hydroxysteroid dehydrogenase/isomerase | 29.92 |
| PFL_6133 |  | NAD dependent epimerase/dehydratase | 29.37 |
| PFL_4307 | WbpV | UDP-Glc 4-epimerase WbpV | 29.41 |
| PFL_3045 | arnA | bifunctional UDP-glucuronic acid decarboxylase/UDP-4-amino-4-deoxy-L-arabinose formyltransferase | 25.98 |
| PFL_4587 |  | NAD dependent epimerase/dehydratase | 34.09 |
| PFL_4375 |  | NAD dependent epimerase/dehydratase | 28.78 |
| PFL_5491 | Gmd | GDP-mannose 4,6-dehydratase | 28.07 |
| PFL_5106 | WbjB | trifunctional UDP-D-GlcNAc 4,6-dehydratase/5-epimerase/3-epimerase WbjB | 24.46 |
| PFL_3633 |  | NAD dependent epimerase/dehydratase | 22.34 |

<sup>a</sup>Predicted function based on Pfam analysis (53).

**Figure 3:**
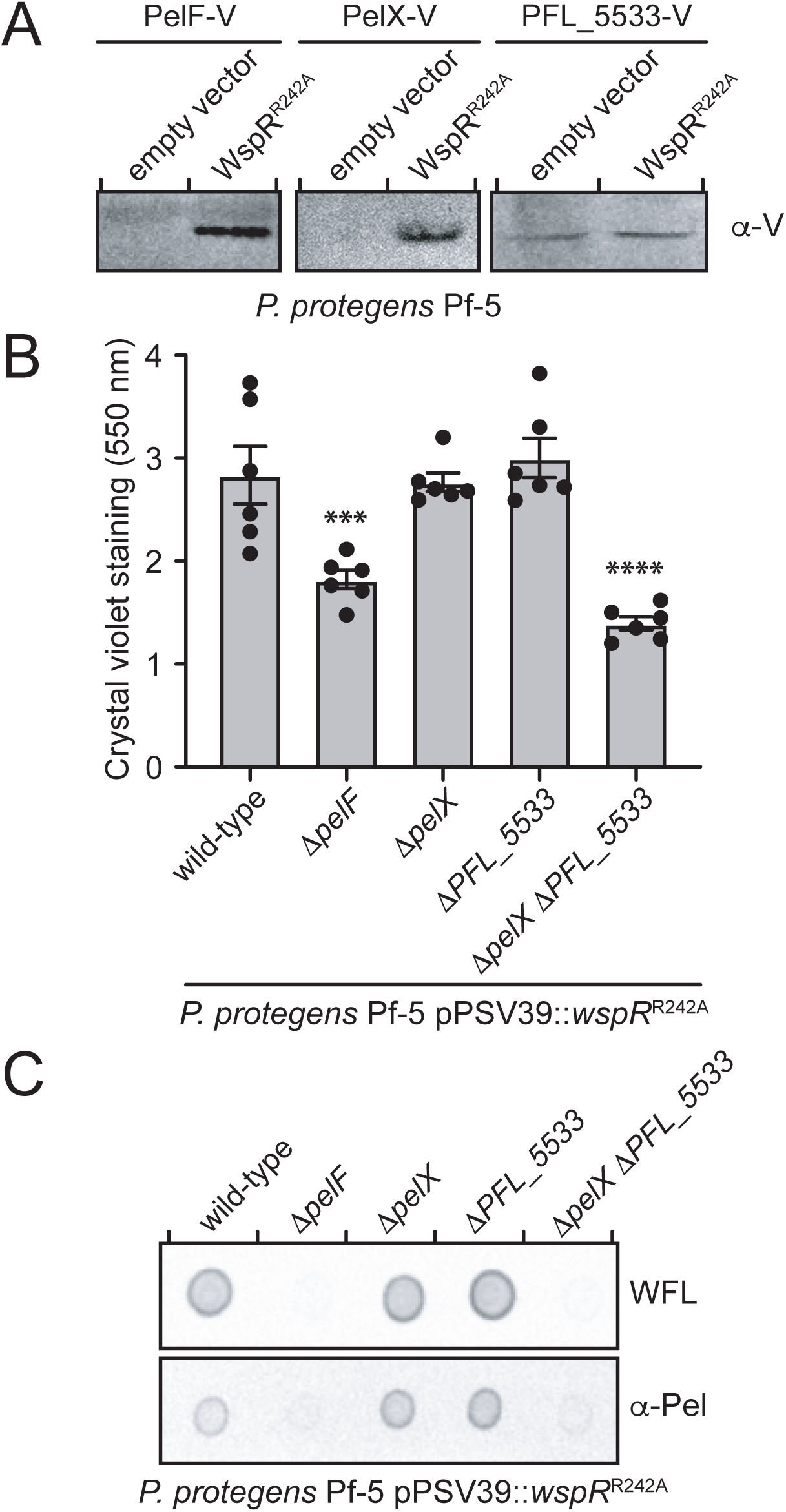
*pelX* or PFL_5533 is required for Pel polysaccharide-dependent biofilm formation by *P. protegens*. (A) Western blot of VSV-G tagged proteins PelF, PelX, or PFL_5533 in the presence of native c-di-GMP levels (empty vector) or elevated c-di-GMP levels (WspR^R242A^). (B) Biofilm biomass as determined using the crystal violet assay after 6 h of static growth. Error bars indicate the standard error of the mean of two independent trials performed in triplicate. Statistical significance was evaluated using one-way ANOVA with Bonferroni correction; ***P < 0.001; ****P < 0.0001. (C) Dot blot of culture supernatants of the indicated strains probed using either horseradish peroxidase conjugated *Wisteria floribunda* lectin (WFL) or the Pel antibody (α-Pel) (10,27).

To determine whether PFL_5533 contributes to biofilm formation by *P. protegens*, we generated a strain lacking this gene and examined biofilm formation in our WspR^R242A^ overexpression background. Similar to our Δ*pelX* strain, we detected no significant difference in biofilm formation between ΔPFL_5533 and wild-type strains **(Fig. 3B)**. In contrast, a Δ*pelX* ΔPFL_5533 double mutant exhibited a defect in biofilm formation comparable to that of a Δ*pelF* strain, which is incapable of producing Pel **(Fig. 3B).** To confirm that this reduction in biofilm formation was due to decreased Pel polysaccharide secretion, *P. protegens* culture supernatants were analyzed using a lectin from *Wisteria floribunda* (WFL) that specifically recognizes terminal GalNAc moieties, and Pel-specific antisera generated using *P. aeruginosa* Pel polysaccharide (10,27). Culture supernatants from wild-type *P. protegens* displayed a strong signal when analyzed by both of these detection methods, while a Δ*pelF* strain exhibited no signal, indicating that these tools can be used to monitor Pel polysaccharide produced by this bacterium (**Fig. 3C**; (9)). In line with our biofilm data, Pel was detected in culture supernatants from Δ*pelX* and ΔPFL_5533 strains at levels comparable to wild-type whereas no Pel polysaccharide was detected in the Δ*pelX* ΔPFL_5533 double mutant. Taken together, these data indicate that *pelX* and PFL_5533 have genetically redundant functions in biofilm formation under our experimental conditions, and that the activity of a predicted SDR family enzyme is essential for Pel polysaccharide biosynthesis and Pel-dependent biofilm formation by *P. protegens*.

### *PelX is a UDP-GlcNAC C4-epimerase that preferentially epimerizes* N*-acetylated UDP-hexoses*

To gain further insight into PelX function, we initiated structural and functional studies on recombinant PelX protein. Initial efforts to purify His_6_-tagged PelX overexpressed in *E. coli* yielded two species consistent with a monomer and dimer of PelX when analyzed by SDS-PAGE. Addition of reducing agent significantly lowered the abundance of the putative PelX dimer, suggesting that this higher molecular weight species likely arose from the formation of an intermolecular disulfide bond. This intermolecular disulfide bond is likely not biologically relevant given that the bacterial cytoplasm is a reducing environment. As sample heterogeneity can be problematic for both the interpretation of biochemical data and protein crystallization, we generated a PelX variant in which the cysteine residue presumed to be involved in disulfide bond formation (C232) was mutated to serine (PelX^C232S^). This PelX^C232S^ variant appeared as a monomer on SDS-PAGE and its purification to homogeneity was straightforward. When examined by size exclusion chromatography, PelX^C232S^ had an apparent molecular weight of 64 kDa compared to its expected monomeric molecular weight of 35 kDa, suggesting that like other characterized SDR enzymes, PelX forms non-covalent, SDS-sensitive dimers in solution **(Fig. S1**; (28)).

The SDR superfamily of enzymes are known to catalyze numerous chemical reactions including dehydration, reduction, isomerization, epimerization, dehalogenation, and decarboxylation (14). We hypothesized that PelX likely functions as an epimerase because UDP-GalNAc, the putative precursor for Pel, is typically generated from UDP-GlcNAc by SDR epimerase-catalyzed stereochemical inversion at the C4 position of the hexose ring. Characterized SDR C4-epimerases are classified into three groups based on their substrate preference (29). Group 1 epimerases preferentially interconvert non-acetylated UDP-hexoses, group 2 epimerases are equally able to interconvert non-acetylated and *N*-acetylated UDP-hexoses, while group 3 epimerases preferentially interconvert *N*-acetylated-UDP-hexoses. Given that the Pel polysaccharide is GalNAc rich, we hypothesized that PelX likely functions as either a group 2 or group 3 epimerase. To examine the potential epimerase activity of PelX, we used ^1^H NMR to monitor the stereochemistry of UDP-GlcNAc, UDP-GalNAc, UDP-Glc, or UDP-galactose (UDP-Gal) in the presence or absence of purified PelX^C232S^. Two ^1^H NMR resonances with characteristic multiplicities in the 5.4-5.7 ppm H-1^”^ region allow for the differentiation of UDP-GalNAc/UDP-Gal from UDP-GlcNAc/UDP-Glc, respectively **(Fig. 4A and 4B).** Using these resonances, we found that PelX^C232S^ readily converts UDP-GalNAc to UDP-GlcNAc and vice versa **(Fig. 4A and 4C)**. PelX^C232S^ also converted a minor amount of UDP-Gal to UDP-Glc, however, we did not observe significant conversion of UDP-Glc to UDP-Gal **(Fig. 4B)**. Collectively, these data define PelX as a group 3 UDP-hexose C4-epimerase.

**Figure 4:**
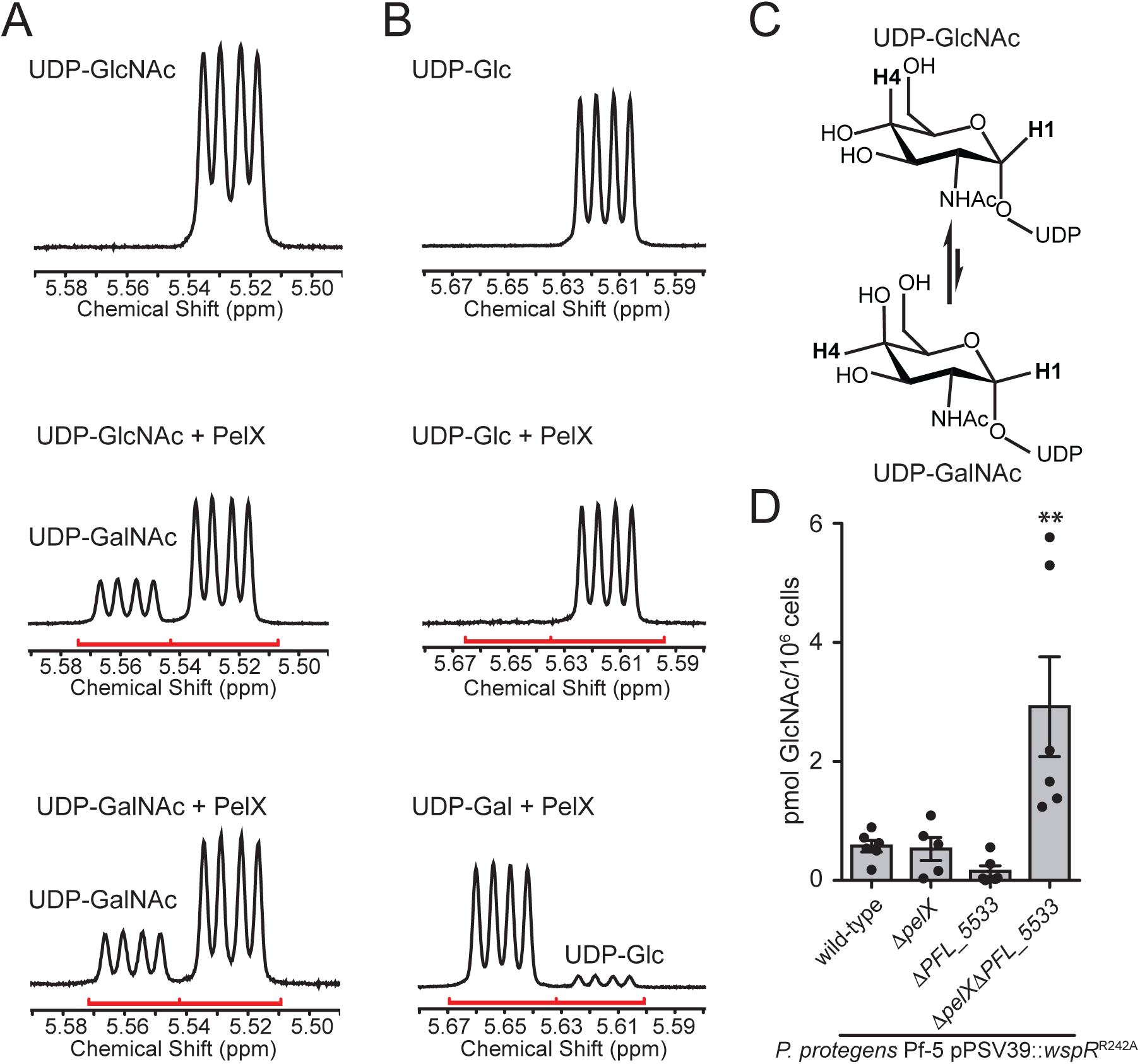
PelX is a Class 3 UDP-GlcNAc C4-epimerase. ^1^H NMR focused on the anomeric H-1 region of the spectra (red line) for (A) UDP-GlcNAc incubated without enzyme (top), UDP-GlcNAc with PelX (middle), and UDP-GalNAc with PelX (bottom) and (B) UDP-Glc incubated without enzyme (top), UDP-Glc with PelX (middle), and UDP-Gal with PelX (bottom). (C) PelX catalyzes the epimerization of UDP-GlcNAc to UDP-GalNAc by inversion of the hydroxyl group at position C4. (D) LC-MS/MS quantification of GlcNAc from cell extracts of the indicated strains. Error bars represent the standard error of the mean of six independent biological replicates. Statistical significance was evaluated using unpaired t-test. *P < 0.05.

To corroborate our biochemical data, we next performed absolute quantification of cellular GalNAc and GlcNAc levels in our WspR^R242A^-expressing *P. protegens* wild-type, Δ*pelX*, ΔPFL_5533, and Δ*pelX* ΔPFL_5533 strains. While GalNAc levels were below the limit of our detection methods, we found that GlcNAc levels were significantly elevated in the epimerase deficient background compared to both wild-type and the individual epimerase mutant strains **(Fig. 4D)**. Taken together with our ^1^H NMR results, these data suggest that PelX and its homologue PFL_5533 function to generate pools of UDP-GalNAc precursors for polymerization into Pel polysaccharide.

### PelX resembles members of the SDR enzyme superfamily

Having established that PelX is a UDP-GlcNAc C4-epimerase, we next sought to determine its structure to obtain further insight into substrate recognition by this enzyme. Despite its straightforward purification and homogenous oligomeric state, we found PelX^C232S^ to be recalcitrant to crystallization. We next attempted to crystallize PelX^C232S^ in complex with its confirmed substrate UDP-GlcNAc. Crystals of PelX^C232S^ incubated with UDP-GlcNAc appeared within three days and the structure of the complex was solved to 2.1 Å resolution using molecular replacement with the SDR family member WbpP (PDB ID: 1SB8) as the search model (28). PelX crystallized in space group *P*2_1_2_1_2 and contains a dimer in the asymmetric unit, an arrangement observed for many other structurally characterized SDR family members (**Fig. 5A**; (30)). The dimer interface of PelX^C232S^ is similar to that observed in the WbpP crystal structure where each protomer contributes two α-helices to a four-helix bundle.

**Figure 5:**
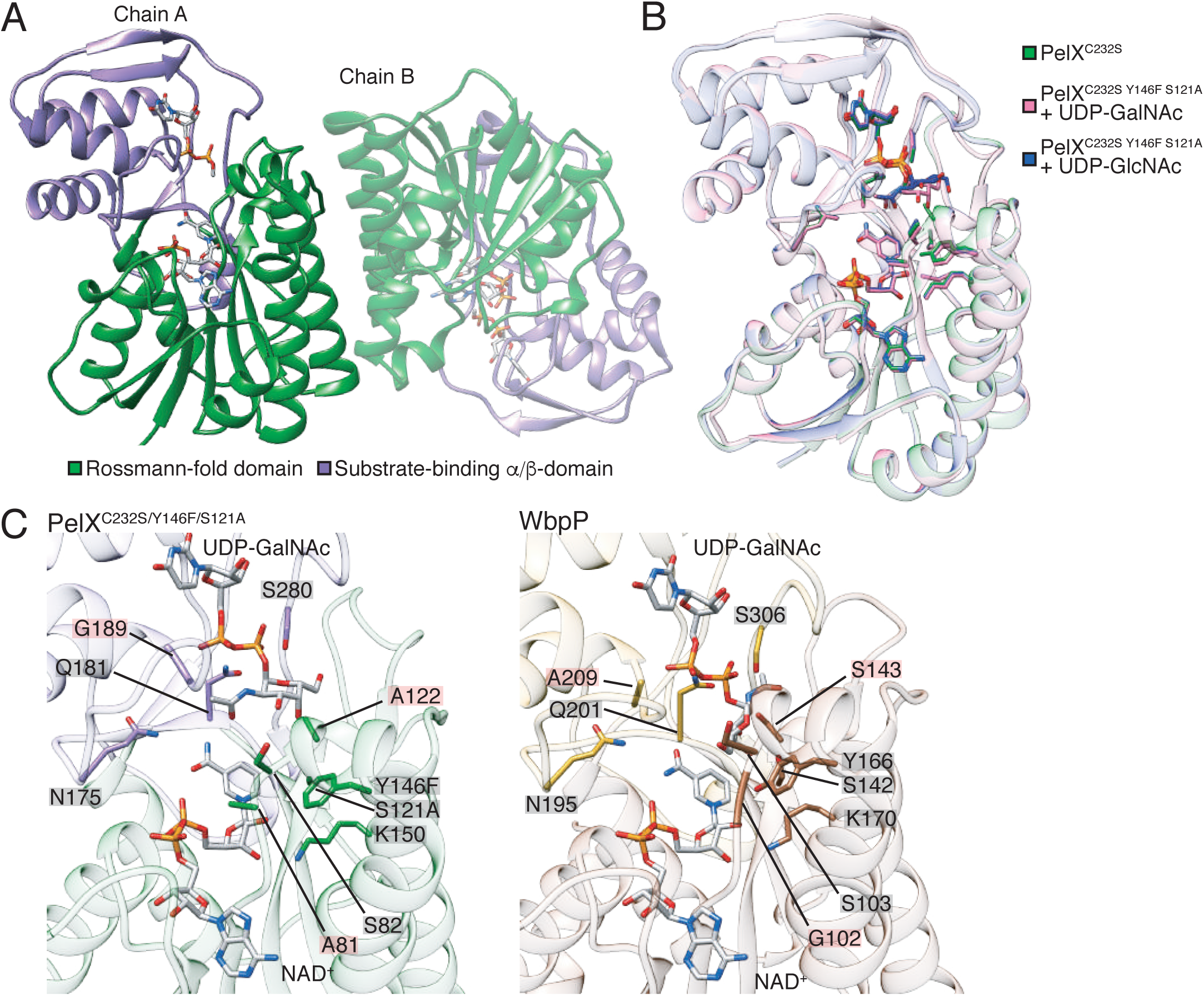
PelX is a member of the short-chain dehydrogenase/reductase superfamily of enzymes. Cartoon representations of PelX^C232S^. (A) PelX^C232S^ is displayed as found in the asymmetric unit with its N-terminal Rossmann-fold domain shown in green, and its C-terminal substrate-binding α/β-domain in purple. (B) Overlay of PelX^C232S/Y146F/S121A^-substrate complexes with the wild-type PelX^C232S^ coloured according to the legend in the figure (C) Comparison of the active site of PelX^C232S/Y146F/S121A^ -UDP-GalNAc complex, and WbpP-UDP-GalNAc complex (PDB ID: 1SB8). UDP-GalNAc and nicotinamide adenine dinucleotide (NAD^+^), and active site residues are shown in stick representation.

The overall structure of PelX^C232S^ shows that it possesses the characteristic domains associated with the SDR family, which includes an N-terminal NAD^+^-binding Rossmann-fold (residues 1-172 and 218-243) and a C-terminal α/β-domain involved in substrate-binding (residues 173-217 and 244-310; **Fig. 5A**). PelX^C232S^ contains the GxxGxxG motif required for binding NAD^+^ that is found in all SDR family members as well as the active site catalytic triad Sx_24_Yx_3_K (31). Although NAD^+^ was not exogenously supplied in the purification or crystallization buffers, electron density for this cofactor was clearly observed, suggesting it was acquired during PelX^C232S^ overexpression in *E. coli*. While the addition of UDP-GlcNAc was essential for the formation of crystals, we were unable to model the GlcNAc moiety of this molecule due to the poor quality of the electron density **(Fig. S3)**. We speculate that the sugar moiety may be disordered because PelX^C232S^ is catalytically active and converting a portion of the UDP-GlcNAc to UDP-GalNAc. Modeling UDP alone rather than UDP-GlcNAc improved the refinement statistics of the overall model and resulted in ligand B-factors comparable to the surrounding protein atoms **(Table 2).**

**Table 2:** Data collection and refinement statistics.

|  | PelX<sup>C232S</sup> | PelX<sup>C232S Y146F S121A</sup> | PelX<sup>C232S Y146F S121A</sup> |
| --- | --- | --- | --- |
|  | + UDP-GlcNAc | + UDP-GlcNAc | + UDP-GalNAc |
| **Data Collection** |  |  |  |
| Wavelength (Å) | 1.075 | 1.075 | 1.075 |
| Space group | *P*2<sub>1</sub>2<sub>1</sub>2 | *P*2<sub>1</sub>2<sub>1</sub>2 | *P*2<sub>1</sub>2<sub>1</sub>2 |
| Cell dimensions |  |  |  |
| *a*, *b*, *c* (Å) | 123.0, 75.5, 79.2 | 124.2, 75.5, 79.3 | 123.6, 75.3, 79.3 |
| $\alpha$, $\beta$, $\gamma$ (°) | 90.0, 90.0, 90.0 | 90.0, 90.0, 90.0 | 90.0, 90.0, 90.0 |
| Resolution (Å) | 50.00 – 2.1 (2.18–2.10) | 50.00 – 2.1 (2.18–2.10) | 50.00 – 2.1 (2.18–2.10) |
| Total no. of reflections | 676002 | 598870 | 589555 |
| Total no. of unique reflections | 43495 | 43637 | 44099 |
| *R*<sub>merge</sub> (%)<sup>b</sup> | 8.0 (58.0) | 12.2 (72.2) | 10.6 (78.7) |
| *I*/σ*I* | 38.8 (5.3) | 23.6 (3.6) | 26.5 (3.1) |
| Completeness (%) | 100.0 (100.0) | 100.0 (100.0) | 99.5 (98.6) |
| Redundancy | 11.4 (14.6)<sup>a</sup> | 7.6 (12.5) | 7.6 (12.2) |
| **Refinement** |  |  |  |
| *R*<sub>work</sub> / *R*<sub>free</sub> (%)<sup>c</sup> | 16.7/19.7 | 15.6/19.5 | 15.6/19.5 |
| Average B-factors (Å<sup>2</sup>) |  |  |  |
| Protein | 41.29 | 31.23 | 35.80 |
| NAD | 32.25 | 21.64 | 25.42 |
| UDP-sugar | 40.82 | 33.13 | 33.19 |
| Water | 41.52 | 37.69 | 40.39 |
| Rms deviations |  |  |  |
| Bond lengths (Å) | 0.006 | 0.006 | 0.006 |
| Bond angles (°) | 0.881 | 0.863 | 0.897 |
| Ramachandran plot (%)<sup>d</sup> |  |  |  |
| Total favored | 97.87 | 97.55 | 97.71 |
| Total allowed | 2.13 | 2.45 | 2.13 |
| Coordinate error (Å)<sup>e</sup> | 0.21 | 0.2 | 0.22 |
| PDB code | 6WJB | 6WJ9 | 6WJA |

<sup>a</sup>Values in parentheses correspond to the highest resolution shell.
<sup>b</sup> $R_{\text{merge}} = \sum |I(k) - \langle I \rangle| / \sum I(k)$ where *I*(*k*) and $\langle I \rangle$ represent the diffraction intensity values of the individual measurements and the corresponding mean values. The summation is over all unique measurements.
<sup>c</sup> $R_{\text{work}} = \sum ||F_{\text{obs}}| - k|F_{\text{calc}}|| / |F_{\text{obs}}|$ where *F*<sub>obs</sub> and *F*<sub>calc</sub> are the observed and calculated structure factors, respectively.
*R*<sub>free</sub> is the sum extended over a subset of reflections (5%) excluded from all stages of the refinement.
<sup>d</sup>As calculated using MOLPROBITY (54).
<sup>e</sup>Maximum-Likelihood Based Coordinate Error, as determined by PHENIX (52).

Previous studies on a catalytically inactive variant of the UDP-Gal 4-epimerase GalE from *E. coli* allowed for the co-crystallization and modeling of UDP-Glc and UDP-Gal in the active site of this enzyme (32). In their study, these authors targeted the serine and tyrosine residues of the consensus Sx_24_Yx_3_K active site motif. Guided by this approach, we generated a variant of PelX^C232S^ with S121A and Y146F mutations and confirmed that this variant is catalytically inactive **(Fig. S2).** PelX^C232S/S121A/Y146F^ crystallized readily with either UDP-GlcNAc or UDP-GalNAc, and both structures were solved to a resolution of 2.1 Å using molecular replacement **(Table 2)**. The final models of PelX^C232S/S121A/Y146F^ in complex with UDP-GlcNAc or UDP-GalNAc were both refined to an R_work_/R_free_ of 15.6%/19.5% **(Table 2).** In these structures, the electron density for the sugar moieties was well defined compared to the PelX^C232S^–UDP-GlcNAc co-crystal structure and allowed for the unambiguous modelling of the expected sugar-nucleotides **(Fig. S3)**. Given that both structures showed improved ligand density for their respective substrates, these structures substantiate our biochemical data showing that UDP-GlcNAc and UDP-GalNAc are substrates for PelX. Examination of the active site of our PelX^C232S/S121A/Y146F^-substrate complexes did not show any significant differences in the positions of active site residues suggesting that both sugar-nucleotides are recognized by the enzyme in a similar manner (**Fig. 5B**). We next compared our substrate-bound PelX^C232S/S121A/Y146F^ structures to the UDP-GalNAc bound structure of the aforementioned UDP-hexose C4-epimerase WbpP from *P. aeruginosa*. WbpP shares 32% sequence identity to PelX and also catalyzes the epimerization of UDP-GlcNAc to UDP-GalNAc (28). The overall structure of WbpP is highly similar to PelX^C232S/S121A/Y146F^ (PDB code 1SB8, rms deviation 1.9 Å over 306 Cα) except that WbpP possesses an additional *N*-terminal α-helix not found in PelX. The active site residues identified as being important for sugar-nucleotide interaction in WbpP are invariant in PelX (**Fig. 5C**) with the exception of A81, A122 and G189 in PelX, which correspond to residues G102, S143 and A209 in WbpP, respectively (28). These differences are not predicted to impair specificity towards the UDP-GlcNAc/GalNAc substrate. Rather, Demendi et al found that bulkier residues (G102K, A209N), and mutation of S143A actually displayed enhanced specificity towards acetylated substrates (33). However, while the positions of the PelX^C232S/S121A/Y146F^ and WbpP active site residues and NAD^+^ cofactor are highly similar, comparison of the bound UDP-GalNAc substrate between the two structures reveals distinct differences in the conformations of the GalNAc moiety (**Fig. 5C**). We suspect that this difference in conformation may be a result of the co-crystallization of UDP-GalNAc with wild-type WbpP whereas to observe electron density for the GalNAc moiety of UDP-GalNAc in complex with PelX we had to mutate two active site residues, S121A and Y146F. The residues equivalent to S121 and Y146 in WbpP make contact with the C4 hydroxyl group of GalNAc and thus are likely involved in substrate orientation. These observations suggest that the conformation of UDP-GalNAc in our mutant PelX co-crystal structure may not represent a state adopted during catalysis, but demonstrate a high degree of conformational freedom of the sugar moiety within the relatively large substrate binding pocket. The GlcNAc moiety of UDP-GlcNAc in our PelX^C232S/S121A/Y146F^-UDP-GlcNAc co-crystal structure was also found in a similar orientation as in our UDP-GalNAc-containing structure. Taking these considerations into account and given that WbpP and PelX share a high degree of sequence similarity and interconvert identical substrates with similar preference, we speculate that the epimerization of *N*-acetylated UDP-hexoses by PelX most likely occurs via a similar catalytic mechanism as proposed for WbpP (28,33). In sum, our structural data support our biochemical studies showing that PelX belongs to the group 3 family of UDP-*N*-acetylated hexose C4-epimerases.

## DISCUSSION

In this study, we report the characterization of the Pel polysaccharide precursor-generating enzyme PelX. Using *P. protegens* Pf-5 as a model bacterium, we found that *pelX* is required for Pel polysaccharide-dependent biofilm formation in a strain that also lacks the *pelX* paralogue, PFL_5533. Guided by our ^1^H NMR analyses and multiple crystal structures, we have shown that PelX functions as a UDP-GlcNAc C4-epimerase and that it preferentially interconverts UDP-GlcNAc/UDP-GalNAc over UDP-Glc/UDP-Gal, defining it as a group 3 UDP-*N*-acetylhexose C4-epimerase. Based on these observations and the data presented herein we propose naming PFL_5533 <u>p</u>olysaccharide UDP-<u>G</u>lc<u>N</u>Ac <u>e</u>pimerase (*pgnE*).

Functional redundancy of sugar-nucleotide synthesizing enzymes in biofilm producing bacteria is not unprecedented. For example, in *P. aeruginosa* PAO1, PslB and WbpW both catalyze the synthesis of GDP-mannose, a precursor molecule required for Psl polysaccharide and A-band lipopolysaccharide (LPS). Like PelX and PgnE, PslB and WbpW have been shown to be genetically redundant as a defect in Psl polysaccharide or A-band LPS is only observed when both *pslB* and *wbpW* are deleted (22). Although *P. aeruginosa* PAO1 has another paralogue of PslB and WbpW, AlgA, the *algD* promoter responsible for transcription of the *algA* gene is not significantly activated in non-mucoid strains such as PAO1 (34). Psl biosynthesis, like Pel, is also regulated by c-di-GMP through FleQ (23) whereas being an integral component of the *P. aeruginosa* outer membrane, the genes responsible for A-band LPS synthesis are constitutively expressed (35). Although, at present what additional glycans PgnE may be involved in producing is unknown, it is clear that the existence of paralogous sugar-nucleotide synthesizing enzymes may be a means of keeping up with metabolic demand during the synthesis of multiple cell surface polysaccharides.

We previously reported the isolation of Pel polysaccharide from *P. aeruginosa* PAO1 and carbohydrate composition analyses showed that it is rich in GalNAc (9). Therefore, the co-regulation of a UDP-GlcNAc C4-epimerase with the *pel* genes likely ensures that adequate quantities of UDP-GalNAc are available for Pel biosynthesis when a biofilm mode of growth is favoured. In contrast to *P. protegens* Pf-5, *P. aeruginosa* PAO1 does not contain a *pelX* gene in its Pel biosynthetic gene cluster, yet this bacterium is also capable of producing Pel polysaccharide (36). In the PAO1 genome, the poorly characterized PA4068 gene is found in the same genomic context as *pgnE* whereby both genes are part of a two-gene operon, with the second gene predicted to encode a dTDP-4-dehydrorhamnose reductase (PA4069/PFL_5534; (37)). In addition, the protein encoded by PA4068 shares 76% identity with PgnE, suggesting that this gene may function analogously to *pgnE* and by extension *pelX*. A ΔPA4068 mutant was found to display a surface attachment defect during secretin induced stress suggesting a role for this gene in surface glycan production (37). However, it has been established that Psl is the primary polysaccharide required for *P. aeruginosa* PAO1 biofilm formation even though this strain is genetically capable of synthesizing Pel (36). Consequently, studies characterizing Pel polysaccharide production by PAO1 have relied on an engineered strain that lacks the ability to produce Psl and expresses the *pel* genes from an arabinose-inducible promoter. It may be that only low levels of UDP-GalNAc are required to sustain Pel polysaccharide production by wild-type PAO1 and thus a second UDP-GlcNAc C4-epimerase that is dedicated to Pel production is not required. In contrast, Pel polysaccharide appears to be a major biofilm matrix constituent in *P. protegens* Pf-5 and thus the higher levels of Pel production in this organism may necessitate the need for increased synthesis of UDP-GalNAc precursors.

The epimerization of UDP-Gal to UDP-Glc by PelX occurs much less efficiently than its *N*-acetylated counterpart. Creuzenet and colleagues noted a similar trend for WbpP, a UDP-GlcNAc C4-epimerase involved in *P. aeruginosa* PAK O-antigen biosynthesis, and hypothesized that the poor efficiency displayed by this enzyme towards non-acetylated substrates means that this reaction is unlikely to occur *in vivo* (38). The equilibrium of the PelX catalyzed epimerization between UDP-GalNAc and UDP-GlcNAc *in vitro* is skewed towards the more thermodynamically stable UDP-GlcNAc epimer. A similar balance for this equilibrium has been documented for other epimerases (38,39). We speculate that the continuous polymerization of UDP-GalNAc by the putative Pel polysaccharide polymerase, PelF, would keep the cellular concentration of UDP-GalNAc low and thus drive the equilibrium towards its production.

In conclusion, this work demonstrates the involvement of a Pel polysaccharide precursor generating enzyme required for biofilm formation in *P. protegens*. Our data linking the production of UDP-GalNAc to Pel polysaccharide production lends genetic and biochemical support to the chemical analyses that showed Pel is a GalNAc-rich carbohydrate polymer (9). Furthermore, the identification of a new Pel polysaccharide-dependent biofilm forming bacterium provides an additional model system that can be used for the characterization of this understudied polysaccharide secretion apparatus.

## EXPERIMENTAL PROCEDURES

### Bacterial strains, microbiological media and physiological buffers

All bacterial strains and plasmids used in this study are listed in Table S1. Jensen’s medium contained per liter of MilliQ water: 5 g NaCl, 2.51 g K_2_HPO_4_, 13.46 g glutamic acid, 2.81 g L-valine, 1.32 g L-phenylalanine, 0.33 g/L MgSO_4_•7H_2_O, 21 mg CaCl_2_•2H_2_O, 1.1 mg FeSO_4_•7H_2_O, 2.4 mg ZnSO_4_•7H_2_O, and 1.25% D-glucose. Semi-solid agar medium in Petri dishes was prepared by adding 1.0% noble agar to Jensen’s medium. A 10 × solution of phosphate buffered saline (PBS) was purchased from Amresco, and diluted, as required in sterile MilliQ water. King’s B medium contained per liter of MilliQ water: 10 g proteose peptone #2 (DIFCO), 1.5 g anhydrous K_2_HPO_4_, 15 g glycerol, and 5 mL MgSO_4._ Lysogeny broth (LB) contained per liter of MilliQ water: 10 g tryptone, 10 g NaCl, and 5 g yeast extract. *E. coli* strains were grown with shaking at 37 °C. *P. protegens* strains were grown at 30 °C. The following concentration of antibiotics were used: gentamicin (Gent) 15 μg ml^-1^ (*E. coli*); Gent 30 μg ml^-1^ (*P. protegens*); kanamycin (Kan), 25 μg ml^-1^. Plasmids were maintained in DH5a(λpir).

### *Bioinformatic identification of PelX among* pel *gene clusters in sequenced bacterial genomes*

We have previously constructed a database of genomes containing *pel* gene clusters using the Geneious platform (12,13,40). Briefly, identification of *pel* gene clusters was made via BLASTP (18) searching of the National Center for Biotechnology Information (NCBI), *Pseudomonas* (41), and *Burkholderia* (42) databases (as of May 6, 2018) using *P. aeruginosa* PAO1 PelC (NP_251752.1) as the query sequence. Annotated genomes encoding PelC orthologs were downloaded from the databases and manually binned according to synteny of the *pel* operon. Conserved domains encoded by open reading frames (ORFs) linked to *pel* loci were queried by searching the Conserved Domain Database (CDD)(43). Visualizations of *pel* gene clusters were drawn to scale using Geneious Prime 2020 and Adobe Illustrator.

### Sequence analysis of PelX and PgnE orthologues

According to the *Pseudomonas* Genome database, PFL_2971 and PFL_5533 belong to the *Pseudomonas* ortholog groups (POG) 020331 and POG001617, respectively. Prior to this study, the POGs were unnamed, therefore based on our observations we have named these POGs as *pelX* and *pgnE*. PelX primary amino acid sequences were aligned using MUSCLE (44) to identify highly conserved amino acid residues. Additionally, the *P. protegens* PelX sequence was submitted to Phyre^2^ to determine the predicted fold of the protein (19). The PelX and PgnE protein sequences from *P. protegens* Pf-5 were obtained from the *Pseudomonas* Genome Database (41). Comparison of the PelX structure to previously determined structures was performed using the DALI pairwise comparison server (45).

### *Construction of* P. protegens *chromosomal mutations*

In-frame, unmarked *pslA* (PFL_4208), *pelF* (PFL_2977), *pelX* (PFL_2971), and PFL_5533 gene deletions in *P. protegens* Pf-5 were constructed using an established allelic replacement strategy (46). Flanking upstream and downstream regions of the open reading frames (ORFs) were amplified and joined by splicing-by-overlap extension PCR (primers are listed in **Table S1**). The *pslA, pelF*, and *pelX*, alleles were generated using forward upstream and downstream reverse primers tailed with *EcoRI* and *XbaI*, restrictions sites, respectively (**Table S1**). The PFL_5533 allele was generated using forward upstream and downstream reverse primers tailed with *EcoRI* and *HindIII* restriction sites, respectively (**Table S1**). This PCR product was gel purified, digested and ligated into pEXG2, and the resulting constructs, pLSM33, pLSM34, and pLSM35, pLSM36 were identified and sequenced as described above.

The VSV-G tagged *pelF, pelX*, and PFL_5533 constructs were generated by amplifying flanking upstream and downstream regions surrounding the stop codon of the ORFs of each gene. The reverse upstream and forward downstream primers (**Table S1)** were tailed with complementary sequences encoding the VSV-G peptide immediately before the stop codon. Amplified upstream and downstream fragments were joined by splicing-by-overlap extension PCR using forward upstream and reverse downstream primers tailed with *EcoRI* and *HindIII* restriction sites, respectively (**Table S1**). These PCR products were gel purified, digested, and ligated into pEXG2, as described above. Clones with positive inserts were verified by Sanger sequencing to generate pLSM37, pLSM38, and pLSM39.

The aforementioned pEXG2 based plasmids were introduced into *P. protegens* Pf-5 via biparental mating with donor strain *E. coli* SM10 (47). Merodiploids were selected on LB containing 60 µg mL^-1^ Gentamicin (Gen) and 25 µg mL^-1^ Irgasan (Irg). SacB-mediated counter-selection was carried out by selecting for double-crossover mutations on no salt lysogeny broth (NSLB) agar containing 5% (w/v) sucrose. Unmarked gene deletions were identified by PCR with primers targeting the outside, flanking regions of *pslA, pelF, pelX*, and PFL_5533 (**Table S1, S2**). These PCR products were Sanger sequenced using the same primers to confirm the correct deletion.

### Generation of WspR overexpression strains

The *wspR* nucleotide sequence from *P. aeruginosa* PAO1 was obtained from the *Pseudomonas* Genome Database and used to design primers specific to full-length *wspR* **(Table S1)**. The forward primer encodes an *EcoRI* restriction site and a ribosomal binding site, while the reverse primer encodes a *HindIII* restriction site. The amplified PCR products were digested with EcoRI and HindIII restriction endonucleases and subsequently cloned into the pPSV39 vector (**Table S1)**. Confirmation of the correct nucleotide sequence of *wspR* was achieved through DNA sequencing (The Center for Applied Genomics, The Hospital for Sick Children). R242 was mutated to an alanine to prevent allosteric inhibition of WspR using the QuickChange Lightning Site Directed Mutagenesis kit (Agilent technologies), as described previously. The resulting expression vector (pLSM-*wspR*^*R242A*^) encodes residues 1-347 of WspR. Introduction of the pPSV39 empty vector or pSLM-*wspR*^*R242A*^ into *P. protegens* was carried out by electroporation. Positive clones were selected for on LB agar containing 30 µg mL^-1^ Gen.

### Crystal violet assay

Overnight cultures grown in King’s B media (KBM), were diluted to a final OD of 0.005 in 1 mL of KBM in a 24-well VDX plate (Hampton Research) and left undisturbed at 30 °C for 120 h. Non-attached cells were removed and the wells were washed thoroughly with water, and stained with 1.5 mL 0.1% (w/v) crystal violet. After 10 minutes, the wells were washed again and the stain solubilized using 2 mL of 95% (v/v) ethanol for 10 minutes. 200 µL was transferred to a fresh 96-well polypropylene plate (Nunc) and the absorbance measured at 550 nm. For strains containing empty pPSV39 or pLSM-*wspR*^*R242A*^, the above protocol was modified slightly. As c-di-GMP significantly upregulated biofilm formation, crystal violet staining for these strains was performed as described previously using 96-well polypropylene plates that were incubated statically for 6 h or 24 h at 30 °C. All strains were grown in KBM containing 30 µg ml^-1^ Gen and 30 µM IPTG.

### Dot blots

Pel antisera was obtained as described in Colvin *et al.* from *P. aeruginosa* PA14 P_BAD_*pel* (10). The adsorption reaction was conducted as described by Jennings *et al* (9). Culture supernatants containing secreted Pel were harvested by centrifugation (16,000 × *g* for 2 min) from 1 mL aliquots of *P. protegens* grown overnight at 30 °C in LB containing 30 µg ml^-1^ Gen and 30 µM IPTG, and treated with proteinase K (final concentration, 0.5 mg mL^-1^) for 60 min at 60 °C, followed by 30 min at 80 °C to inactivate proteinase K.

Pel immunoblots were performed as described by Colvin *et al* (10) and Jennings *et al* (9). 5 μL of secreted Pel, prepared as described above, was pipetted onto a nitrocellulose membrane and left to air dry for 10 min. The membrane was blocked with 5% (w/v) skim milk in Tris-buffered saline (10 mM Tris-HCl pH 7.5, 150 mM NaCl) containing 0.1% (v/v) Tween-20 (TBS-T) for 1 h at room temperature and probed with adsorbed α-Pel at a 1:60 dilution in 1% (w/v) skim milk in TBS-T overnight at 4 °C with shaking. Blots were washed three times for 5 min each with TBS-T, probed with goat α-rabbit HRP-conjugated secondary antibody (Bio-Rad) at a 1:2000 dilution in TBS-T for 45 min at room temperature with shaking, and washed again. All immunoblots were developed using SuperSignal West Pico (Thermo Scientific) following the manufacturer’s recommendations.

For WFL-HRP immunoblots, 5 μL of secreted Pel, prepared as described above, was pipetted onto a nitrocellulose membrane and left to air dry for 10 min. The membrane was blocked with 5% (w/v) bovine serum albumin (BSA) in TBS-T for 1 h at room temperature and probed with 10 μg/mL of WFL-HRP (EY Laboratories) in 2% (w/v) BSA in TBS-T with 0.2 g/L CaCl_2_ overnight at room temperature with shaking. Membranes were washed twice for 5 min and once for 10 min with TBS-T, then developed as described above.

### Western blot sample preparation and analysis

For analysis of protein levels from WspR^R242A^ overexpressing strains containing VSV-G-tagged PelF, PelX or PFL_5533, 5 mL of LB media containing 30 μM IPTG and 30 μg mL^-1^ Gen was inoculated with the appropriate strain and allowed to grow overnight at 30 °C with shaking. Culture density was normalized to an OD_600_ = 1 and 1 mL of cells was centrifuged at 5,000 × g for 5 min to pellet cells. The cell pellet was resuspended in 100 µL of 2× Laemmli buffer, boiled for 10 min at 95 °C, and analyzed by SDS-PAGE followed by Western blot. For Western blot analysis, a 0.2 µm polyvinylidene difluoride (PVDF) membrane was wetted in methanol and soaked for 5 min in Western transfer buffer (25 mM Tris-HCl, 150 mM glycine, 20% (v/v) methanol) along with the SDS-PAGE gel to be analyzed. Protein was transferred from the SDS-PAGE gel to the PVDF membrane by wet transfer (25 mV, 2 h). The membrane was briefly washed in TBS-T before blocking in 5% (w/v) skim milk powder in TBS-T for 2 h at room temperature with gentle agitation. The membrane was briefly washed again in TBS-T before incubation overnight with α-VSV-G antibody in TBS-T with 1% (w/v) skim milk powder at 4 °C. The next day, the membrane was washed four times in TBS-T for 5 min each before incubation for 1 h with secondary antibody (1:2000 dilution of BioRad affinity purified mouse α-rabbit IgG conjugated to alkaline phosphatase) in TBS-T with 1% (w/v) skim milk powder. The membrane was then washed three times with TBS-T for 5 min each before development with 5-bromo-4-chloro-3-indolyl phosphate/nitro blue tetrazolium chloride (BioShop ready-to-use BCIP/NBT solution). Developed blots were imaged using a BioRad ChemiDoc imaging system.

### Cloning and mutagenesis

The *pelX* nucleotide sequence from *P. protegens* Pf-5 (PFL_2971) was obtained from the *Pseudomonas* Genome Database (41) and used to design primers specific to full-length *pelX* (**Table S1**). The amplified PCR products were digested with *Nde*I and *Xho*I restriction endonucleases and subsequently cloned into the pET28a vector (Novagen). Confirmation of the correct nucleotide sequence of *pelX* was achieved through DNA sequencing (ACGT DNA Technologies Corporation). The resulting expression vector (pLSM-PelX) encodes residues 1-309 of PelX fused to a cleavable N-terminal His_6_ tag (His_6_-PelX) for purification purposes (**Table S2**). To prevent aggregation of PelX in solution, a non-conserved cysteine (C232) was mutated to a serine with the aid of the QuickChange Lightning Site Directed Mutagenesis kit (Agilent technologies) and confirmed with DNA sequencing (ACGT DNA Technologies Corporation). The PelX^C232S^ active site mutant (S121A Y146F) was generated analogously.

### Expression and purification of PelX

The expression of PelX^C232S^ was achieved through the transformation of the PelX^C232S^ expression vector into *Escherichia coli* BL21 (DE3) competent cells, which were then grown in 2 L lysogeny broth (LB) containing 50 µg mL^-1^ kanamycin at 37 °C. The cells were grown to an OD_600_ of 0.6 whereupon isopropyl-β-D-1-thiogalactopyranoside (IPTG) was added to a final concentration of 1.0 mM to induce expression. The induced cells were incubated for 20 h at 25 °C prior to being harvested *via* centrifugation at 6 260 × *g* for 20 min at 4 °C. The resulting cell pellet was stored at -20 °C until required.

The cell pellet from 2 L of bacterial culture was thawed and resuspended in 80 mL of Buffer A [50 mM Tris-HCl pH 8.0, 300 mM NaCl, 5% (v/v) glycerol, and 1 mM tris(2-carboxyethyl)phosphine (TCEP)] containing 1 SIGMAFAST protease inhibitor EDTA-free cocktail tablet (Sigma). Due to the presence of two remaining cysteines in PelX^C232S^, TCEP was included to prevent intermolecular cross-linking of the protein. These cysteines are not predicted to be involved in disulfide bond formation given their poor conservation and the cytoplasmic localization of PelX^C232S^. The resuspension was then lysed by homogenization using an Emulsiflex-C3 (Avestin, Inc.) at a pressure between 15 000 – 20 000 psi, until the resuspension appeared translucent. Insoluble cell lysate was removed by centrifugation for 25 min at 25 000 × *g* at 4 °C. The supernatant was loaded onto a 5 mL Ni^2+^-NTA column pre-equilibrated with Buffer A containing 5 mM imidazole in order to reduce background binding. To remove contaminating proteins, the column was washed with 5 column volumes of Buffer A containing 20 mM imidazole. Bound protein was eluted from the column with 3 column volumes of Buffer A containing 250 mM imidazole. SDS-PAGE analysis revealed that the resulting His_6_-PelX^C232S^ was ∼99% pure and appeared at its expected molecular weight of 35 kDa. Fractions containing PelX^C232S^ were pooled and concentrated to a volume of 2 mL by centrifugation at 2 200 × *g* at a temperature of 4 °C using an Amicon ultra centrifugal filter device (Millipore) with a 10 kDa molecular weight cut-off. PelX^C232S^ was purified and buffer exchanged into Buffer B [20 mM Tris-HCl pH 8.0, 150 mM NaCl, 5% (v/v) glycerol, 1 mM TCEP] by size-exclusion chromatography using a HiLoad 16/60 Superdex 200 gel-filtration column (GE Healthcare). PelX^C232S^ eluted as a single Gaussian shaped peak, and all PelX^C232S^ containing fractions were pooled and concentrated by centrifugation, as above, to 8 mg mL^-1^ and stored at 4 °C. PelX^C232S/Y146F/S121A^ was purified similarly.

### Determination of the PelX oligomerization state by gel filtration analysis

Oligomerization of PelX^C232S^ was determined using a Superdex 200 10/300 GL column (GE Life Sciences). The column was equilibrated in Buffer B. Molecular weight standards (Sigma, 12-200 kDa) were applied to the column as directed. PelX^C232S^ was applied to the column at 7.5 mg ml^-1^ (100 µL) and protein elution was monitored at 280 nm.

### NMR activity assay

The following method has been adapted from Wyszynski et al (39). Enzymatic reactions were performed in 30 mM sodium phosphate pH 8.0, with 50 µg of freshly purified PelX^C232S^ and 10 mM UDP-GlcNAc, UDP-Glc, UDP-Gal, or 5 mM UDP-GalNAc in a total reaction volume of 220 µL. After incubation at 37 °C for 1 hour, the mixture was flash frozen and lyophilized. The resulting material was dissolved in 220 µL of D_2_O and analyzed by ^1^H NMR. As control experiments, the same procedures were applied to samples lacking PelX or UDP-GlcNAc. Data were collected on a Varian 600 MHz NMR spectrometer.

### Intracellular metabolite extraction

*P. protegens* Pf-5 wild-type, Δ*pelX*, ΔPFL_5533 and Δ*pelX* ΔPFL_5533 strains that had been transformed with a plasmid expressing WspR^R242A^ (pLSM21) were streaked out twice in succession on Jensen’s agar containing 30 µg/mL gentamicin, and these first and second subcultures were grown for 48 h at 30 °C. For each biological replicate, cells from the second subcultures were collected using a polyester swab and suspended in 30 mL of Jensen’s medium to match an optical density at 600 nm (OD_600_) of 0.6. Subsequently, two 10 mL aliquots of this standardized culture were each passed through a syringe filter (0.45 µm, PVDF, Millipore) to collect the bacteria. These filters were placed face-up on Jensen’s agar using flame sterilized tweezers and were then incubated at 30 °C for 3 h.

Following this incubation, the first filter was placed in 2 mL of sterile PBS containing 1 mM purified recombinant PelA, and this filter was incubated for 30 min at room temperature to break up aggregates (11). An established microtiter dilution method for viable cell counting was used to determine the number of bacteria on the filter (48). The second filter was put into a Petri dish (60 x 15 mm) containing 2 mL of cold 80% (v/v) LC-MS grade methanol, which was incubated for 15 min. Afterwards, 1 mL of 80% LC-MS grade methanol was used to wash the filter, and then the 3 mL of the methanol extract were transferred to a 5 mL microcentrifuge tube. These tubes were placed in a centrifuge at 7000 × g for 30 min at 4 °C, and then 2 mL of the supernatant were transferred to a 2 mL microcentrifuge tube. The methanol was evaporated using a speed vac, and then the dried cell extracts were suspended in 200 µL of 50% (v/v) LC-MS grade methanol. Cell extracts were then stored at -80 °C until LC-MS analysis. Mass spectrometry measurements of GalNAc were normalized to viable cell counts.

### Liquid chromatography-mass spectrometry (LC-MS)

Mass spectral data collection was done on a Thermo Scientific Q Exactive™ Hybrid Quadrupole-Orbitrap™ Mass Spectrometer in negative ion full scan mode (70-1000 m/z) at 140,000 resolution, with an automatic gain control target of 1e^6^, and a maximum injection time of 200 ms. A Thermo Scientific Ion Max-S API source outfitted with a HESI-II probe was used to couple the mass spectrometer to a Thermo Scientific Vanquish Flex UHPLC platform. Heated electrospray source parameters for negative mode were as follows: spray voltage -2500 V, sheath gas 25 (arbitrary units), auxiliary gas 10 (arbitrary units), sweep gas 2 (arbitrary units), capillary temperature 275 °C, auxiliary gas temperature 325 °C. A binary solvent mixture of 20 mM ammonium formate at pH 3.0 in LC-MS grade water (solvent A) and 0.1% (v/v) formic acid in LC-MS grade acetonitrile (solvent B) were used in conjunction with a Syncronis™ HILIC LC column (Thermo Fisher Scientific 97502-102130) to achieve chromatographic separation of chemical compounds. For these runs the following gradient was used at a flow rate of 600 uL min^-1^: 0-2 min, 100% B; 2-15 min, 100-80% B; 15-16 min, 80-5% B; 16-17.5 min, 5% B; 17.5-18 min, 5-100% B; 18-21 min, 100 % B. For all runs the sample injection volume was 2 uL. Raw data acquisition was carried out using Thermo Xcalibur 4.0.27.19 software. Data analysis was carried out using MAVEN software (49). Compound identification was achieved through matching of high-resolution accurate mass and retention time characteristics to those of authentic standards. Secondary compound confirmation was performed by matching of fragmentation profiles obtained through parallel reaction monitoring.

### Crystallization and structure determination

Commercial sparse matrix crystal screens from Microlytic (MCSG1-4) were prepared at room temperature (22 °C) with PelX^C232S^ at a concentration of 8 mg mL^-1^ (0.23 mM). UDP-GlcNAc was added exogenously to a concentration of 2 mM. Trials were set up in 48-well VDX plates (Hampton Research) by hand with 3 µL drops at a ratio of 1:1 protein to crystallization solution over a reservoir containing 200 µL of the crystallization solution. Crystal trays were stored at 22 °C. The best crystals were obtained from condition 32 [0.2 M ammonium sulphate, 0.1 M sodium citrate pH 5.6, 25% (w/v) PEG 4000] from MCSG-1 (Microlytic). This condition yielded stacked flat square plate crystals that took approximately 5 days to grow to maximum dimensions of 300 µm x 300 µm x 50 µm. PelX was unable to form crystals in the absence of UDP-GlcNAc.

Crystals of PelX^C232S^ were cryoprotected in well solution supplemented with 20% (v/v) ethylene glycol by briefly soaking the crystal in a separate drop. Crystals were soaked for 2-3 s prior to vitrification in liquid nitrogen, and subsequently stored until X-ray diffraction data were collected on beamline X29A at the National Synchrotron Light Source (NSLS) at Brookhaven National Laboratory. A total of 360 images of 1° Δφ oscillation were collected on an ADSC Q315 CCD detector with a 250 mm crystal-to-detector distance and an exposure time of 0.4 s per image. The data were processed using DENZO and integrated intensities were scaled using SCALEPACK from the HKL-2000 program package (50). The data collection statistics are summarized in **Table 2**. The structure was solved by molecular replacement using WbpP as a model with PHENIX AutoMR wizard. The resulting map was of good quality and allowed manual model building using COOT (51,52). The model was then refined using PHENIX.REFINE (52) to a final R_work_/R_free_ of 16.7% and 19.7%, respectively.

PelX^C232S/S121A/Y146F^ in complex with UDP-GalNAc or UDP-GlcNAc was crystallized under the same conditions as the wild-type protein, and data collection and refinement were performed as described above. The corresponding statistics can be found in **Table 2**.

## Acknowledgements

This work was supported in part by the grants from the Canadian Institutes of Health Research (CIHR) to PLH (MOP 43998 and FDN154327), the National Institute of Health 2R01AI077628 to MRP, and the Natural Sciences and Engineering Research Council (NSERC) to (JJH 435631-2013). PLH and JJH are recipients of Canada Research Chairs. LSM, GBW, and JCW have been supported by Canada Graduate Scholarships from NSERC. LSM and JCW have been supported by graduate scholarships from the Ontario Graduate Scholarship Program, and The Hospital for Sick Children Foundation Student Scholarship Program. GBW has been supported by a graduate scholarship from Cystic Fibrosis Canada. Crystallization utilized the Structural and Biophysical Core Facility at The Hospital for Children supported in part by the Canadian Foundation for Innovation. Beam line X29 at the National Synchrotron Light Source is supported by the US Department of Energy and the NIH National Center for Research Resources. The coordinates and structure factors for PelX^C232S^ in complex with NAD^+^ and UDP, PelX^C232S S121A Y146F^ UDP-GlcNAc and PelX^C232S S121A Y146F^ UDP-GalNAc have been deposited in the PDB, ID codes 6WJB, 6WJA, 6WJ9, respectively. Metabolomics data were acquired by R.A.G. at the Calgary Metabolomics Research Facility (CMRF), which is supported by the International Microbiome Centre and the Canada Foundation for Innovation (CFI-JELF 34986). I.A.L. is supported by an Alberta Innovates Translational Health Chair.

## Conflict of interest

The authors declare that they have no conflicts of interest with the contents of this article.

## Author contributions

L.S.M., M.R.P., and P.L.H. conceptualization; L.S.M., G.B.W., J.J.H., I.A.L., and P.L.H. formal analysis; L.S.M., G.B.W., H.R., R.P., R.J.W, T.E.R., E.R., A.O., R.A.G., and I.A.L., investigation; L.S.M., G.B.W., and P.L.H. writing-original draft; L.S.M., G.B.W., J.C.W., J.J.H., and P.L.H. writing-review and editing; M.N., H.R., M.R.P., and I.A.L. resources; J.J.H, I.A.L., and P.L.H. supervision; P.L.H. funding acquisition; P.L.H. project administration.

## FIGURE LEGENDS

**Figure S1:**
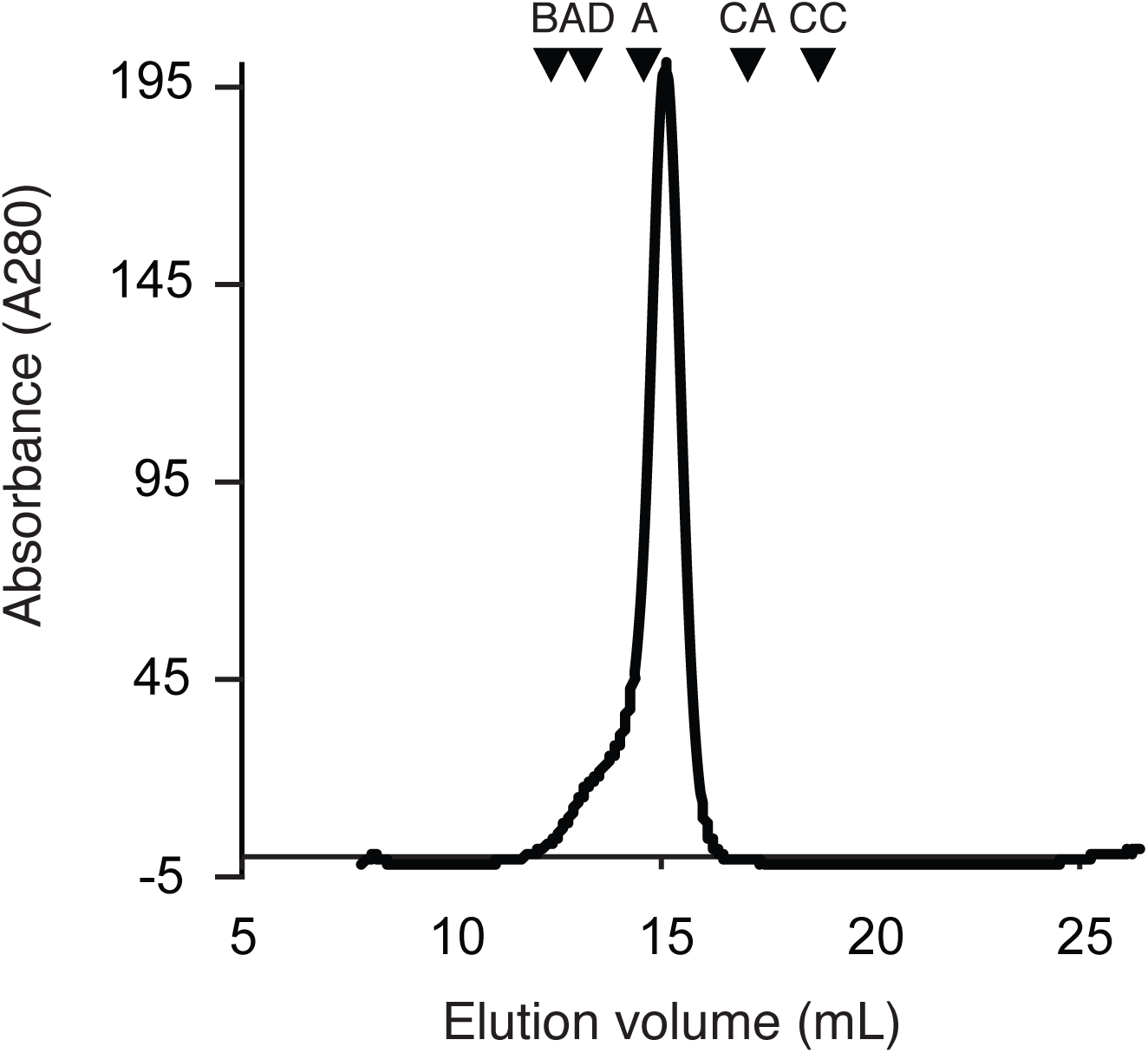
PelX^C232S^ forms a dimer in solution. Analytical gel filtration demonstrates that PelX^C232S^ exists as a dimer in solution, eluting at a molecular weight of approximately 63 kDa. Expected molecular weight: 34.6 kDa. Protein standards used to calibrate the column are indicated by inverted triangles; BA, β-amylase; AD, alcohol dehydrogenase; A, albumin; CA, carbonic anhydrase; CC, cytochrome C. The molecular weights of β-amylase, alcohol dehydrogenase, albumin, carbonic anhydrase, and cytochrome C are 200 kDa, 150 kDa, 66 kDa, 29 kDa, and 12.4 kDa, respectively.

**Figure S2:**
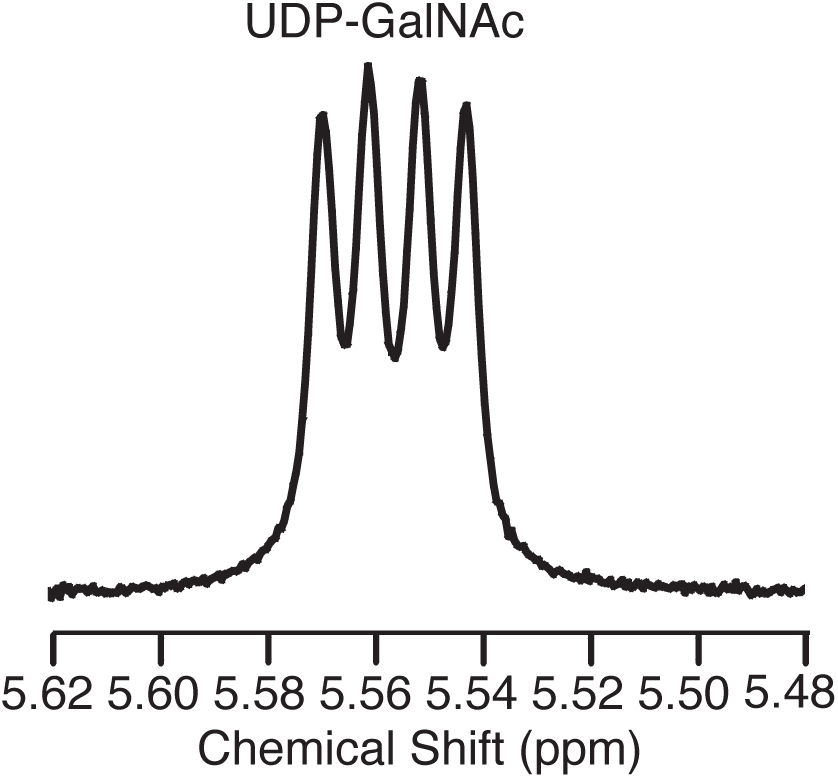
PelX^C232S/Y146F/S121A^ is catalytically inactive towards UDP-GalNAc. ^1^H NMR spectrum from the reaction of PelX^C232S S121A Y146F^ with UDP-GalNAc.

**Figure S3:**
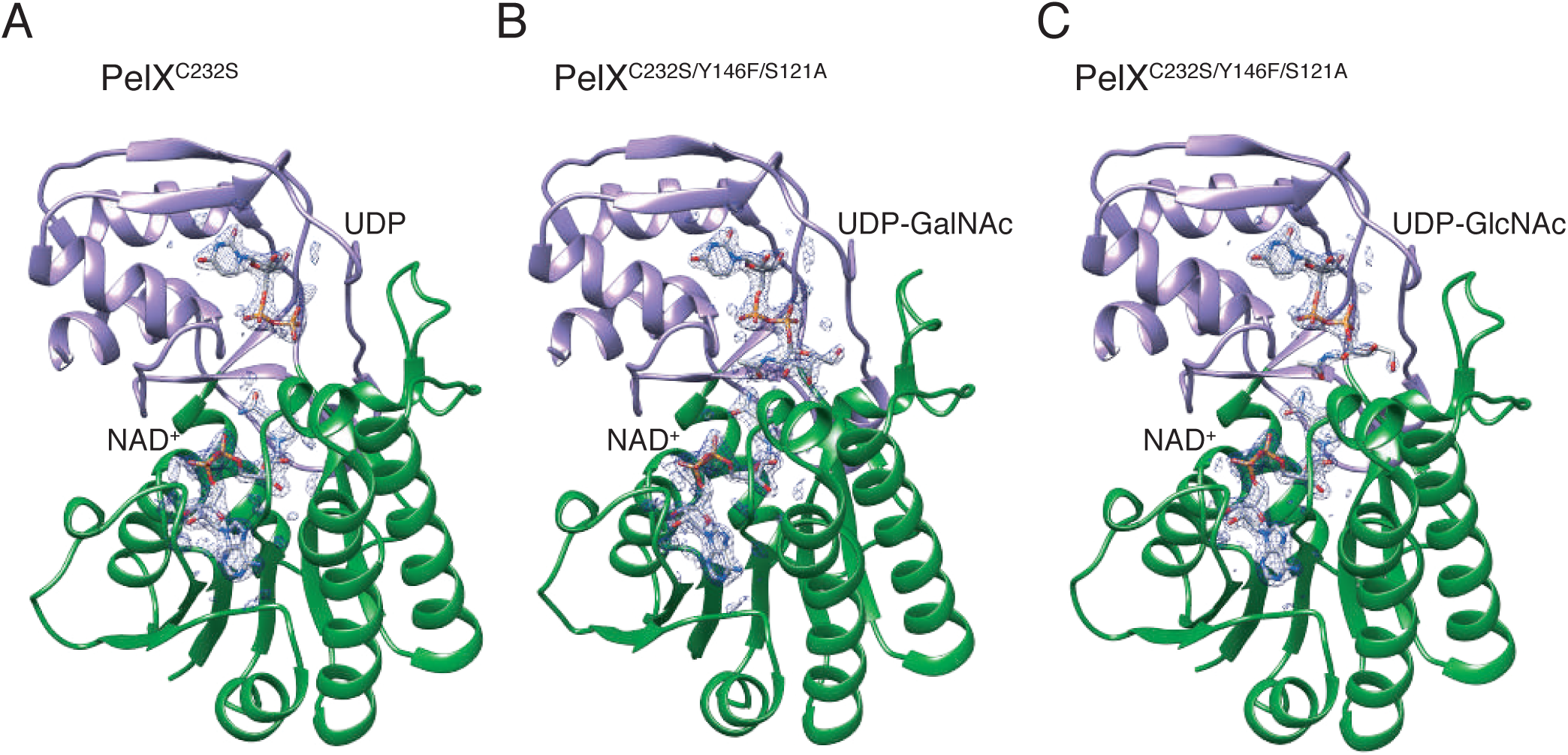
PelX^C232S/Y146F/S121A^ density of the ligands. PelX is displayed with its N-terminal Rossmann-fold domain shown in green, and its C-terminal substrate-binding α/β-domain in purple as in Figure 5. (A) PelX^C232S^ with density shown for UDP and NAD^+^ (B) PelX^C232S/Y146F/S121A^ in complex with UDP-GalNAc and NAD^+^ and (C) PelX^C232S/Y146F/S121A^ in complex with UDP-GlcNAc and NAD^+^. All three structures were modeled with NAD^+^ and nucleotide or sugar-nucleotide shown in stick representation, with the corresponding |2mFo-DFc| map displayed as black mesh contoured at 2.0 σ.

## Notes

### Competing Interest Statement

The authors have declared no competing interest.

