## Supporting Information for "PelX is a UDP-*N*-acetylglucosamine C4-epimerase involved in Pel polysaccharide-dependent biofilm formation"

Running title: *Pel biosynthesis requires a UDP-GlcNAc C4-epimerase*

**Keywords**: polysaccharide, biofilm, *Pseudomonas protegens*, X-ray crystallography, epimerase

**Table S1: Bacterial strains and plasmids used in this study**

| **Strain/Plasmid/Primer** | **Genotype/Properties** | | | **Source** |
| --- | --- | --- | --- | --- |
| ***E. coli*** | | | | |
| TOP10 | F^-^ *mcr*AΔ(*mrr*-*hsd*RMS-*mcr*BC) Φ80*lac*Z ΔM15 Δ*lac*X74 *rec*A1 *ara*D139 Δ(ara leu) 7697 *gal*U *gal*K *rps*L *end*A1 *nup*G, Str^R^ | | | Invitrogen |
| BL21CodonPlus^TM^ (DE3)-RP | F^-^ *ompT* *hsdS*(rB^−^ mB^−^)*dcm*^+^ Tet^r^ *gal*λ (DE3) *end*AHte *met*A∷Tn5(Kan^r^) [*arg*U *ile*Y *leu*W Cam^r^] | | | Stratagene |
| DH5α | F^–^ Φ80*lac*ZΔM15 Δ(*lac*ZYA-*arg*F) U169 *rec*A1 *end*A1 *hsd*R17 (rK–, mK+) *pho*A *sup*E44 λ– *thi*-1 *gyr*A96 *rel*A1 | | | Invitrogen |
| SM10 | *thi thr leu tonA lacY supE recA::RP4-2-Tc::Mu Km λpir*, Kan^R^, Tet^R^ | | | 1 |
| ***P. protegens*** | | | | |
| Pf-5 | Wild-type | | |  |
| Pf-5 Δ*pslA* | In frame deletion of *pslA* (PFL_4208) | | | This study |
| Pf-5 Δ*pelF* | In frame deletion of *pelF* (PFL_2977) | | | This study |
| Pf-5 Δ*pslA* Δ*pelF* | In frame deletion of *pslA* and *pelF* | | | This study |
| Pf-5 Δ*pelX* | In frame deletion of *pelX* (PFL_2971) | | | This study |
| Pf-5 ΔPFL_5533 | In frame deletion of PFL_5533 | | | This study |
| Pf-5 Δ*pelX* ΔPFL_5533 | In frame deletion of *pelX* and PFL_5533 | | | This study |
| Pf-5 *pelX-V* | *pelX* with a C-terminal VSV-G (YTDIEMNRLGK) tag | | | This study |
| Pf-5 *pelF-V* | *pelF* with a C-terminal VSV-G (YTDIEMNRLGK) tag | | | This study |
| Pf-5 PFL_5533*-V* | PFL_5533 with a C-terminal VSV-G (YTDIEMNRLGK) tag | | | This study |
| **Plasmids** | | | | |
| **Protein production** | | | | |
| pET-28a(+) | IPTG inducible expression vector; Kan^r^ | | | Novagen |
| pLSM-PelX_Pp_^1-309^ _C232S_ | PelX_Pp_ encoding the full-length protein. Expressed thrombin-cleavable fusion protein MGSSH_6_SSGLVPRGSHM-PelX_Pp_^1-309^ in pET-28a. Mutation of cysteine 232 to serine. | | | This study |
| pLSM-PelX_Pp_^1-309^ _C232S/Y146F/S121A_ | PelX_Pp_ encoding the full-length protein. Expressed thrombin-cleavable fusion protein MGSSH_6_SSGLVPRGSHM-PelX_Pp_^1-309^ in pET-28a. Mutation of cysteine 232 to serine, tyrosine 146 to phenylalanine, and serine 121 to alanine. | | | This study |
| **WspR overexpression** | | | | |
| pPSV39 | | Expression vector with *lacI*, lacUV5 promoter derived from pPSV35 | 2 | |
| pLSM21 | | WspR^R242A^ from PAO1 encoding the full-length protein with a mutation to the  allosteric inhibition site in pPSV39. IPTG inducible expression. | This study | |
| **Two Step Allelic exchange** | | | | |
| pEXG2 | allelic exchange vector, Gen^R^ | | | 2 |
| pLSM33 | pEXG2 with an in-frame deletion allele for *P. protegens* Pf-5 *pslA* (PFL_4208) Gen^R^ | | | This study |
| pLSM34 | pEXG2 with an in-frame deletion allele for *P. protegens* Pf-5 *pelF* (PFL_2977) Gen^R^ | | | This study |
| pLSM35 | pEXG2 with an in-frame deletion allele for *P. protegens* Pf-5 *pelX* (PFL_2971) Gen^R^ | | | This study |
| pLSM36 | pEXG2 with an in-frame deletion allele for *P. protegens* Pf-5 PFL_5533 Gen^R^ | | | This study |
| pLSM37 | pEXG2 with full-length *pelF* fused to a C-terminal VSV-G tag Gen^R^ | | | This study |
| pLSM38 | pEXG2 with full-length *pelX* fused to a C-terminal VSV-G tag Gen^R^ | | | This study |
| pLSM39 | pEXG2 with full-length PFL_5533 fused to a C-terminal VSV-G tag Gen^R^ | | | This study |

Amp, ampicillin; Cam, chloramphenicol; Kan, kanamycin; Gen, Gentamicin; Str, streptomycin; Tet, tetracycline

**Table S2: Primers used in this study**

| **Primers** | **Sequence*** |
| --- | --- |
| **Vectors for recombinant protein production** | |
| PelX 1F | GG **CAT ATG** TCT GCC GAA CGG ATA CTG |
| PelX 309R | GG **CTC GAG** CTA AAG GCT GCG GTA AAG CCG |
| PelX C232S F | CAG GGC CTG GAA aGC CCC GCG CCG G |
| PelX C232S R | C CGG CGC GGG GCt TTC CAG GCC CTG |
| PelX Y146F F | CC CTC ACG CCC TtC GCG GCG GAC AA |
| PelX Y146F R | TT GTC CGC CGC GaA GGG CGT GAG GG |
| PelX S121A F | G GTG TTC GCG TCC gcT GCG GCG GTC TAT GG |
| PelX S121A R | CC ATA GAC CGC CGC Agc GGA CGC GAA CAC C |
| **Allelic exchange vectors** |  |
| PslA_Pf5_upF | TAG TAC AGA **GAA TTC** GGC AAT CAG CCG GGT ATG G |
| PslA_Pf5_upR | *G TGT GCT CAT TCA GAA GGC CTC* GCT GTC TAC AGG TTG CAA CCG |
| PslA_Pf5_downF | GAG GCC TTC TGA ATG AGC ACA C |
| PslA_Pf5_downR | T CAA TCA GTA **TCT AGA** GCA TGC AGA ACA TCC CGC CG |
| PelF_Pf5_upF | TAG TAC AGA **GAA TTC** CAA CTG CAT TCG CCC ACC TTC |
| PelF_Pf5_upR | *CAC AGC CTC CTT GTG TGG GGT* GGG GGT TGA GTC GGG GGT |
| PelF_Pf5_downF | ACC CCA CAC AAG GAG GCT GTG |
| PelF_Pf5_downR | T CAA TCA GTA **TCT AGA** CAG CGC CAG CAA CAG CCCT |
| PelX_Pf5_upF | TAG TAC AGA **GGT ACC** GGA ACA ACG CATAGG GAA TCGC |
| PelX_Pf5_upR | *GCT GCG GTA AAG CCG TTC CAG* AAC CAG TAT CCG TTC GGC AGA CAT |
| PelX_Pf5_downF | CTG GAA CGG CTT TAC CGC AGC |
| PelX_Pf5_downR | T CAA TCA GTA **AAG CTT** TCG GGG AGC AAC TGG AAA CTG |
| PFL_5533 upF | GTG **GAA TTC** GCT TAC TAC TTC GAC TGG TTT C |
| PFL_5533 upR | *GGC GAG GCC GAC GCT CAT* CAA TAC GAG GCC TTC AGC CAT |
| PFL_5533 downF | ATG AGC GTC GGC CTC GCC |
| PFL_5533 downR | GGT **AAG CTT** CAA CGC CCA CGA AGC TGG T |
| PelX VSVG upF | GGG **GAA TTC** ATG GCA ATA ACG GCG AGG GC |
| PelX VSVG upR | ttt tcc taa tct att cat ttc aat atc tgt ata AAG GCT GCG GTA AAG CCG TTC |
| PelX VSVG downF | tat aca gat att gaa atg aat aga tta gga aaa TAG TTC GTT GCC TTA GGG GCG |
| PelX VSVG downR | GGT **AAG CTT** CTG GTC CTG CAG CGC CTT G |
| PelF VSVG upF | GGG **GAA TTC** GGT GGT GCC GAT CAA GGA CG |
| PelF VSVG upR | ttt tcc taa tct att cat ttc aat atc tgt ata CAC AGC CTC CTT GTG TGG GG |
| PelF VSVG downF | tat aca gat att gaa atg aat aga tta gga aaa TAA ATG GCC GGC ATC GGT TTC G |
| PelF VSVG downR | GGG **AAG CTT** ATC ACC AGG AGG ATG CGG TTA TA |
| PFL_5533 VSVG upF | GGG **GAA TTC** ATG GCA ACA ACG GGG AAG GC |
| PFL_5533 VSVG upR | ttt tcc taa tct att cat ttc aat atc tgt ata GCG CCC CAT CAG GCG GG |
| PFL_5533 VSVG downF | Tat aca gat att gaa atg aat aga tta gga aaa TGA GGG AAA ACA CGC ACA TGA AA |
| PFL_5533 VSVG downR | GAG **AAG CTT** ACC CCG TTC GAA CAC GAC GT |
| **Sequencing Primers** | |
| PslA_Pf5_seqF | GGC TGG CCG GGG CGT C |
| PslA_Pf5_seqR | GGT GGC TCT GCT CCA GGC A |
| PelF_Pf5_seqF | GCG AGG CGC AGA CGT GGC |
| PelF_Pf5_seqR | CAC CAG CTT CTC GGC CCG |
| PelX_Pf5_seqF | TGC AGC AAG GAG GTG CGG G |
| PelX_Pf5_seqR | GGC GAC GCC ATC GAG CTC |
| PFL_5533 seqF | AAC TGC TCG ACG ACA CCC |
| PFL_5533 seqR | GCA CGA ACG ATG ATG TCA C |
| **WspR overexpression** | |
| WspR-F | GC **GAA TTC** AGG AGG ATA TTC ATG CAC AAC CCT CAT GAG AGC |
| WspR-R | GC **AAG CTT** TCA GCC CGC CGG GGC |

*Restriction site sequences are in **bold**; regions of complementary to the target amplicon are underlined; lower case letters denote a nucleotide substitution or mismatch to sequence (in the case of VSV-G tag); regions of complementarity to facilitate splicing of PCR products are in *italics*
